# Metabolic compatibility and the rarity of prokaryote endosymbioses

**DOI:** 10.1101/2022.04.14.488272

**Authors:** Eric Libby, Christopher Kempes, Jordan Okie

## Abstract

The evolution of the mitochondria was a significant event that gave rise to the eukaryotic lineage and most large complex life. Central to the origins of the mitochondria was an endosymbiosis between prokaryotes. Yet, despite the potential benefits that can stem from a prokaryotic endosymbiosis, their modern occurrence is exceptionally rare. While many factors may contribute to their rarity, we lack methods for estimating the extent to which they constrain the appearance of a prokaryotic endosymbiosis. Here, we address this knowledge gap by examining the role of metabolic compatibility between a prokaryotic host and endosymbiont. We use genome-scale metabolic models from three different databases (AGORA, KBase, and CarveMe) to assess the viability, fitness, and adaptability of potential prokaryotic endosymbioses. We find that while more than half of host-endosymbiont pairings are metabolically viable, the resulting endosymbioses have reduced growth rates compared to their ancestral metabolisms and are unlikely to gain mutations to overcome these fitness differences. In spite of these challenges, we do find that they may be more robust in the face of environmental perturbations at least in comparison with the ancestral host metabolism lineages. Our results provide a critical set of null models and expectations for understanding the forces that shape the structure of prokaryotic life.

## Introduction

The evolution of mitochondria—from independent organism to intracellular organelle—is exceptional in terms of its physiological significance, ecological repercussions, and apparent evolutionary rarity [1, 2, 3]. While prokaryotes and eukaryotes of similar cell size have roughly the same total metabolic rate [4, 5], the advantages conferred by the mitochondria, such as extra scale-free internal membrane area [6] along with distributed copies of the metabolic genes [7], may have facilitated the evolution and spread of large cells and complex multicellularity. Moreover, after establishment of the mitochondria, many branches of the eukaryotic lineage acquired other intracellular endosymbionts, including plastids [8], nitrogen-fixing bacteria [9], and methanogenic archaea [10]. The impressive radiation of Eukaryota and the frequency by which the eukaryotic lineage has gained intracellular endosymbioses suggests they confer evolutionary and ecological advantages [11, 12]. Nevertheless, despite any possible advantages, the evolution of mitochondria is one of only a few reported cases of an endosymbiosis between prokaryotes [13, 14, 15], which is surprising given the opportunities afforded by their abundance and long evolutionary history. Why are there so few documented extant examples of prokaryote endosymbioses and why have they not achieved anything comparable to the remarkable ecological and evolutionary success of the eukaryotic lineage?

Historically, these questions have mainly been addressed by identifying possible barriers to the initial establishment of a prokaryotic endosymbiosis, such as the absence of phagocytosis in modern prokaryotes [16] or issues with metabolic compatibility [17]. Yet, the initial establishment is not the only stage at which a nascent endosymbiosis may encounter limitations. For example, a nascent endosymbiosis must be fit enough to compete with other organisms in the environment and it must also be able to adapt in order to spread into other environments and diversify (see Figure 1). At different stages we may expect various ecological, physiological, or evolutionary constraints to be more dominant, but we currently lack quantitative approaches or null models to estimate the magnitude of these constraints or compare their influence. Thus we have a fundamental knowledge gap concerning the forces that shape prokaryotic evolution, the origin of eukaryotes, and the distribution of endosymbioses in the biosphere. To address this knowledge gap, we need quantitative approaches that clarify the relative importance of the various barriers that limit the biosphere’s production of prokaryotic endosymbioses.

**Figure 1:**
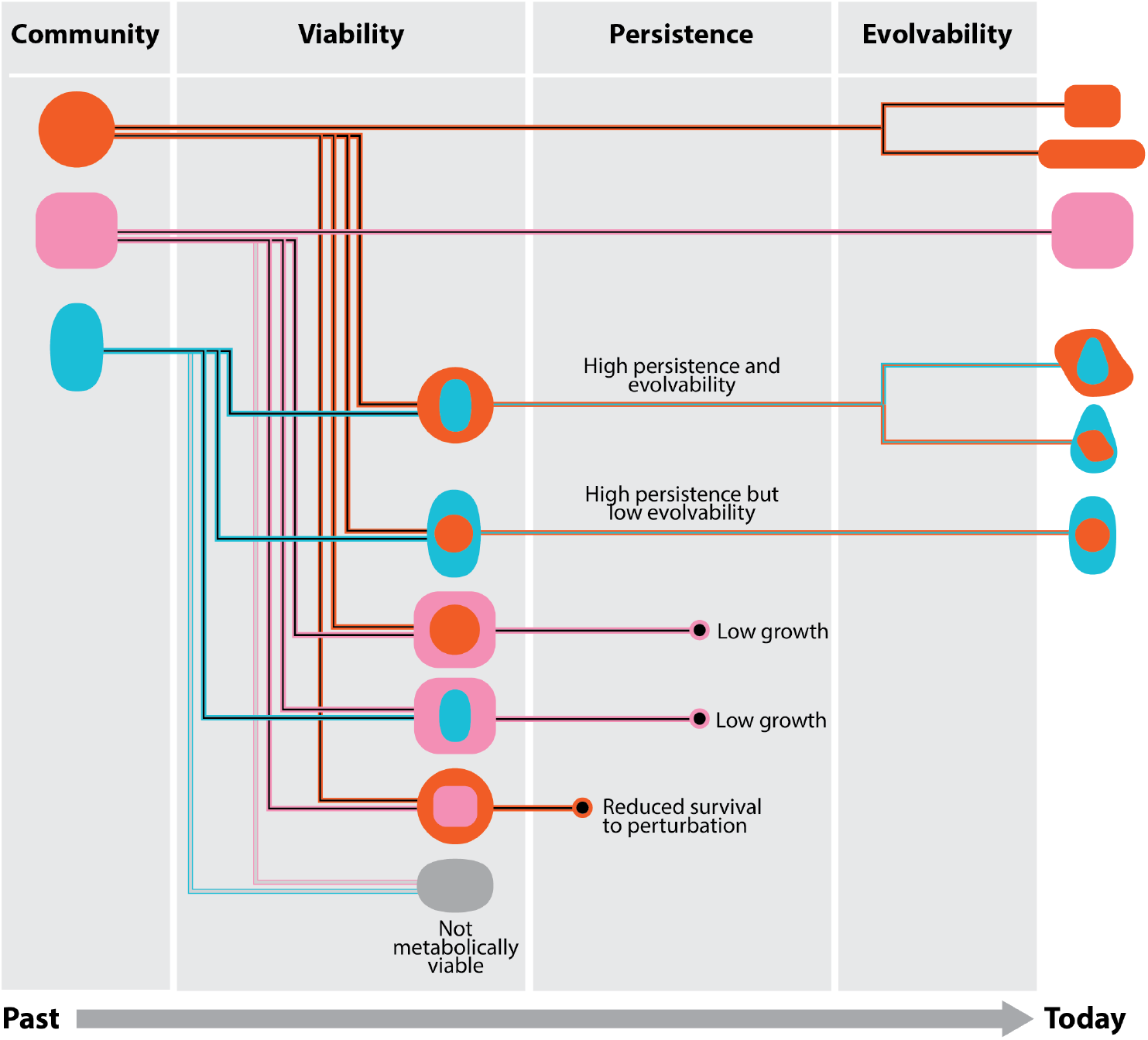
Stages in the evolution of endosymbioses. From its inception a nascent endosymbiosis faces different barriers that challenge the survival and diversification of its lineage. The schematic organizes these barriers into three broad stages corresponding to initial viability, persistence, and evolvability. In each stage metabolic compatibility plays a role in determining which endosymbioses will survive. In the viability stage both host and endosymbiont must be able to grow and reproduce such that the pair can produce offspring host-endosymbiont pairs. In the persistence stage, the endosymbiosis competes with other species including its ancestors. In the evolvability stage, the endosymbiosis must fix beneficial mutations in order to adapt to new environmental conditions. The evolutionary trajectories that successfully pass through all of these stages will determine the abundance and diversity of endosymbioses.

While in principle we could focus on any of the different barriers, one powerful approach is to focus on universal metabolic traits that apply across the biosphere’s ensemble of species and environments [18, 19]. Since metabolism plays a central role in many ecological and evolutionary phenomena [20, 21, 22, 23], a quantitative metabolic framework may offer key insights into the evolution of prokaryotic endosymbioses. Indeed, there are crucial metabolic considerations underlying each stage along the way from initiation to a persistent, adaptable endosymbiosis. For example, the initial viability of an endosymbiont depends on whether it can access all of its required molecular compounds from within the host cell, which is governed by the spatial structure of their coupled metabolisms. Moreover, the fitness and persistence of a host-endosymbiont pair is determined by whether the two metabolisms compete for the same resources or are able to synergize their metabolic pathways; and their adaptability is shaped by the degree to which their coupled metabolisms can harness the effects of mutations. In general it may be difficult to predict precisely the viability, fitness, and adaptability of a putative prokaryotic endosymbiosis because they depend on many idiosyncratic factors, such as the physical and ecological environment, or detailed considerations of which other species co-exist at a particular point in history. However, we can draw upon the burgeoning wealth of genomic data and metabolic network models to elucidate the relative extent to which the metabolic underpinnings of viability, fitness, and adaptability hinder the establishment, evolution, and radiation of prokaryote endosymbioses across the biosphere.

An important tool in quantifying the eco-evolutionary role of metabolism is the genome-scale metabolic model. Such models use the genomes of an organism to infer its metabolic repertoire and predict its growth in different environments. Genome-scale metabolic models are now available for thousands of species and have been successfully applied to a variety of bioengineering, ecological, and evolutionary problems [24, 25]. Of particular relevance, they have also been used to accurately predict fitness and adaptive evolution in prokaryotic communities [26, 27, 28, 29, 30]. In addition to modeling systems that can be experimentally validated, they have also been used to address questions that extend beyond our knowledge of and application to extant life. For example, metabolic models coupled with the network expansion method have been used to propose possibilities for early metabolism on Earth [31], network structure scaling with size [32], and what metabolisms are likely to exist in distinct planetary conditions [33]. Such generality and flexibility is ideal for our interest in exploring the challenges of an event that is possible but rarely seen.

Here we harness the abundance and flexibility of genome-scale metabolic models to consider putative endosymbioses between random pairs of prokaryotes. By using different databases for metabolic models along with a large number of genomes, we are able to quantitatively study general features of metabolic compatibility in nascent endosymbioses and estimate viability, fitness, and adaptability. In this analysis we assess the extent to which metabolic compatibility acts as a barrier, independent of other considerations such as whether endosymbioses form through phagocytosis or via other eco-evolutionary processes [34]. Ultimately we find that the dominant metabolic barriers to a prokaryotic endosymbiosis are due to relative fitness effects and adaptability, not to the initial metabolic viability.

## Results

To assess metabolic compatibility in prokaryotic endosymbioses, we need a broad set of genome-scale metabolic networks (see Methods: Genome-scale metabolic model curation). Thus, we collected metabolic models for prokaryotes from three of the largest databases: AGORA (818 models), KBase (1,637 models), and CarveMe (5,587 models). A common feature across databases is that models organize reactions and compounds into compartments; all models have compartments for the cytoplasm and extracellular environment, but CarveMe models are the only ones with an additional compartment for the periplasm. The databases vary in what species, reactions, and compounds they include, as well as their description of how reactions and compounds behave across compartments. Since the databases also differ in their standards for metabolic model creation and formats, even if the same species appears in multiple databases the metabolic models are not directly translatable. We manage the variation between databases by keeping models from different databases distinct and performing analyses independently on each database. This reduces the risk of introducing errors and provides an opportunity to assess the extent to which our results depend on metabolic model formats.

With our broad collection of metabolic networks, we can assess the metabolic viability of putative prokaryotic endosymbioses. We considered 100,000 random pairs of networks sampled from each database, which were split into 100 sets of 1000 to estimate variation. For each pair of networks, we constructed two metabolic models of the endosymbiosis where networks swapped roles as host and endosymbiont (see Figure 2A). Importantly, every metabolic network in our dataset includes an environment—or equivalently a set of available extracellular compounds—that enable the organism (network) to grow. If we combine the environments from two metabolic networks, we obtain a joint environment in which both networks can grow independently. Thus, we can assess viability of an endosymbiosis by determining whether the host-endosymbiont system can grow in the joint environment (see Methods: Assessing growth and viability of metabolisms).

**Figure 2:**
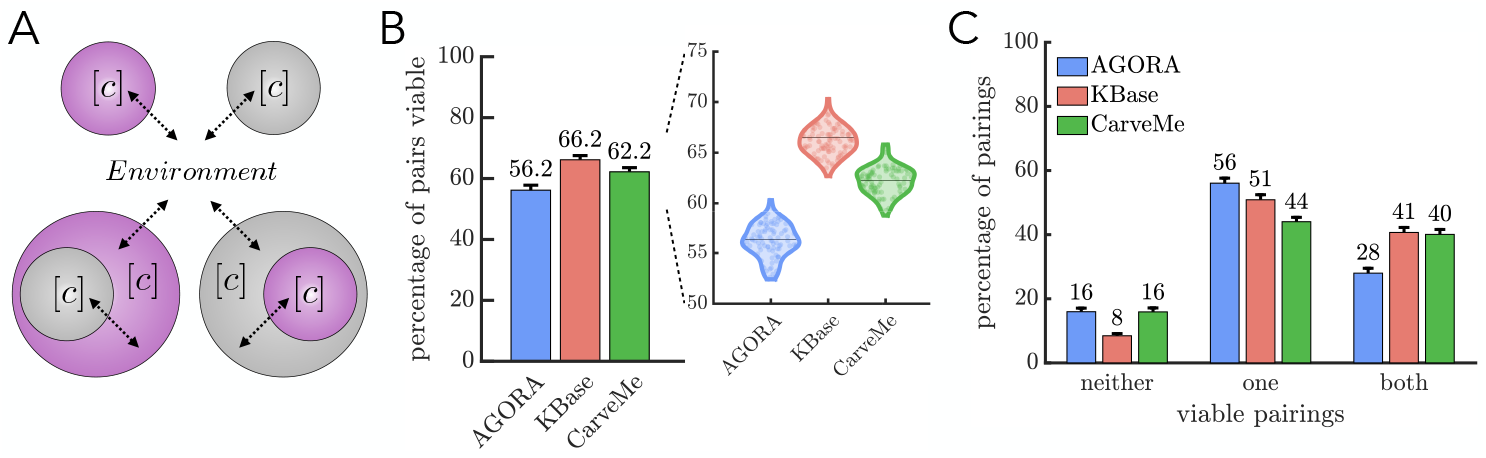
Viability of prokaryotic endosymbioses. a) A schematic shows the nested compartment structure of the two possible host-endosymbiont pairs considered in our analyses. Each cell has a cytoplasm compartment [c] and can exchange compounds with its external environment (indicated by arrows). In endosymbioses the extracellular compartment of the endosymbiont is the cytoplasm compartment of its host. b) A bar graph with an inset of a violin plot shows the percentages of paired metabolisms that form viable endosymbioses for each of the three databases. The percentages are the means of 100 samples of 1000 pairs and are of similar magnitude, 56.2 − 66.2%; however an analysis of variance confirms the means do differ across databases (*p <* .001). c) Bar graphs show the percentage of pairings in which neither, only one, or both configurations of endosymbioses are viable. For all databases there is at least one configuration viable for 84 − 92% of pairs and the most common scenario is where only one configuration is viable.

Figure 2B shows that the average percentage of viable host-endosymbiont pairs varies between 56.2% and 66.2% across the three databases. An analysis of variance confirms that the averages between databases are statistically different (p-value < .001). Such differences between databases may stem from many possible factors including model format, the set of reactions used, the particular compounds available in the environment, and/or the composition of species and their representative environments. We can observe other differences between databases if we evaluate factors that may predict whether an endosymbiosis is viable—such as similarity in reactions or biomass compounds between host and endosymbiont (see Supplementary material: Predicting viability). Yet, despite such differences between databases, all 300 sets of samples have a viable percentage between 52 − 71%, indicating a similar scale for viability estimates. Moreover, if we consider both possible configurations of endosymbioses we find that across databases the majority of pairs, 84 − 92%, have at least one configuration that is viable (see Figure 2C), with most of those pairs (52 − 67%) having just one, specific configuration that is viable.

Within the context of metabolic models, there are two non-exclusive reasons that host-endosymbiont pairs may be nonviable in an environment where each can survive separately (see Figure 3A). First, the host may lack a way of transporting a resource (or waste product) that the endosymbiont requires (or produces) in order to grow. Overcoming this cause of nonviability requires transport of a compound (or compounds) between the host’s cytoplasm and the extracellular environment. Such transport typically occurs in metabolic models via particular reactions, and so the host-endosymbiont pair could be made viable by including the appropriate transport reactions. The second cause of nonviability stems from the endosymbiont needing to perform a reaction whose compounds never enter the cell. Since the endosymbiont cannot directly access the external environment, it cannot perform the necessary reaction. This obstacle can only be overcome if the endosymbiont has direct contact between its membrane and the environment, which is infeasible based on the compartmental structure of an endosymbiosis. Thus, we focused on the first cause of nonviability and evaluated how easily host-endosymbiont pairs can be made viable.

**Figure 3:**
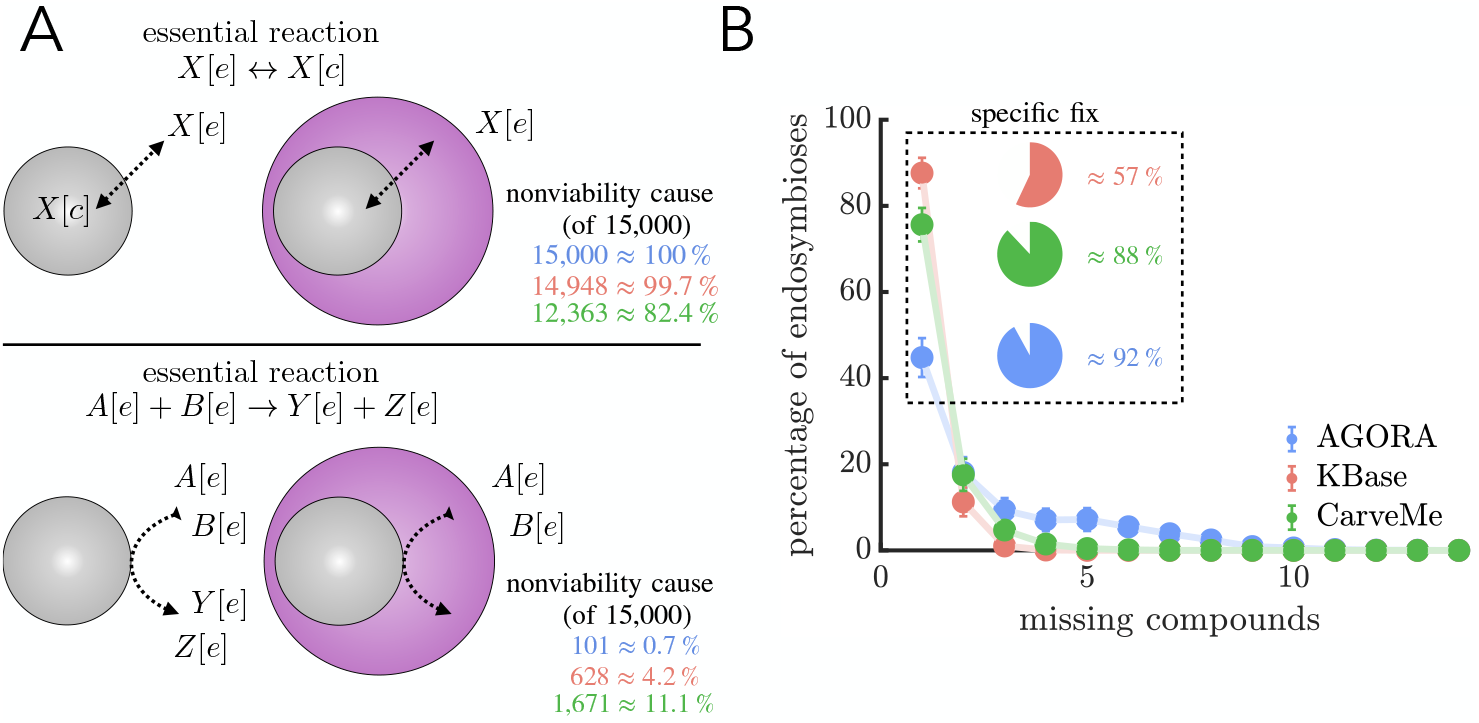
Paths to fixing nonviable endosymbioses. A) A schematic shows the two causes of nonviability in endosymbioses using metabolic models: 1. (top) missing transport of an extracellular compound into the cytoplasm and 2. (bottom) no access to compounds that remain in the extracellular compartment. For each we show a typical form of the missing essential reaction and the percentage of 15,000 nonviable endosymbioses that can be made viable by providing the necessary type of reaction (colors indicate databases, following the legend in B). Between the two causes, the first is more frequently a source of nonviability. We note that within a database the percentages for the two causes do not sum to 100% because they are not mutually exclusive. B) The graph shows the minimum number of compounds whose transport needs to be provided in order to fix nonviable endosymbioses. Each point is the mean of 100 samples of 100 fixable, nonviable endosymbioses and the error bars are the standard deviations of those samples. For each database, the most common case is that the endosymbiosis can be fixed by transporting a single compound. The pie chart inset shows how often that single compound is a specific compound, i.e. there is only one such compound whose transport makes the endosymbiosis viable.

We assessed how often nonviability stems from the host lacking a transport reaction by allowing hosts the ability to transport any compounds that exist in both cytoplasm and extracellular compartments. We attempted to repair 15,000 nonviable endosymbioses in each database and found that the majority can be made viable (100% for AGORA, 99.7% for KBase, and 82.4% for CarveMe). Although adding transport mechanisms for all compounds can fix the majority of nonviable endosymbioses, the likelihood that nonviability can be overcome depends on the number of transport mechanisms needed for viability. We estimated the minimum number of missing transport mechanisms by finding a set of compounds that fix viability only through their combined transport, i.e. failure to transport any one of them leads to nonviability. Using a series of linear programs (see Methods: Fixing viability) on 10,000 randomly sampled nonviable—yet fixable—endosymbioses from each database, we find that the majority of nonviable pairings can be fixed by the addition of a single transport mechanism (Figure 3B). Yet, of those endosymbioses that can be fixed by a single transport mechanism the majority (57 − 92%) require a specific compound without which they cannot be viable (See Figure 3B).

Even if a nascent endosymbiosis is viable it still faces the challenge of competing against a background population composed of its direct ancestors. In principle the competition can take many forms depending on the nature of selection. We first consider competition in terms of the ability of metabolisms to remain viable in the face of environmental degradation. Since host-endosymbiont pairs can draw on two sets of reactions, they may be better able to adjust to changing resource pools so as to have a survival advantage compared to their ancestors. We evaluated this possibility by randomly choosing 10,000 viable host-endosymbiont pairs and determining whether they remain viable when a single compound is removed from their environment. We investigated all possible environments where a single compound is missing, and in each modified environment we also assessed the viability of the ancestral host and endosymbiont metabolisms for comparison.

We observe similar population-level patterns of survival to environmental degradation between the host-endosymbiont pair and its ancestral metabolisms (see Figure 4A and Figure S3). Furthermore we find that the metabolisms act identically in 93 − 97% of perturbations, with the most frequent occurrence being that all survive (see Figure 4B). Of the 3 − 7% of environmental perturbations where we see differences in survival between metabolisms, three scenarios occur most frequently: 1) only the ancestral endosymbiont survives, 2) only the ancestral endosymbiont does not survive, and 3) only the ancestral host does not survive (see Figure 4C). As a consequence, the host-endosymbiont pair survives more environmental perturbations than the ancestral host metabolism in all databases (see Supplementary Figure S4). The outcome of the hostendosymbiont pair versus the ancestral endosymbiont metabolism depends on the database; in KBase models host-endosymbiont pairs survive more often while AGORA and CarveMe show the reverse (see Supplementary Figure S4). In addition to survival selection, genome-scale metabolic models enable us to consider competition in terms of population growth rates. Instead of simply considering a binary outcome of viable versus nonviable, we can use a metabolism’s computed maximum rate of biomass production as a proxy for its populationlevel growth rate. Actual growth rates may depend on many additional factors including gene expression profiles, regulation, and other cell physiological states. To control for some of this variability we compare the endosymbiosis to its constituent metabolisms and avoid comparisons across endosymbioses. Computing the fitness of viable host-endosymbiont pairs, we find that they grow slower than both the ancestral host and endosymbiont metabolisms in over 85% of samples for each of the three databases (see Figures 5A-D). In the rare cases where the host-endosymbiont pair grows faster than its ancestors, its average growth-rate advantage is smaller in magnitude than its average disadvantage when the ancestral metabolisms grow faster. For example, in the AGORA database when the host-endosymbiont pair grows faster than the ancestral host metabolism its mean advantage is 18.9% relative to the host, but when the ancestral host metabolism grows faster its mean advantage is 66.9% relative to the host-endosymbiont pair.

**Figure 4:**
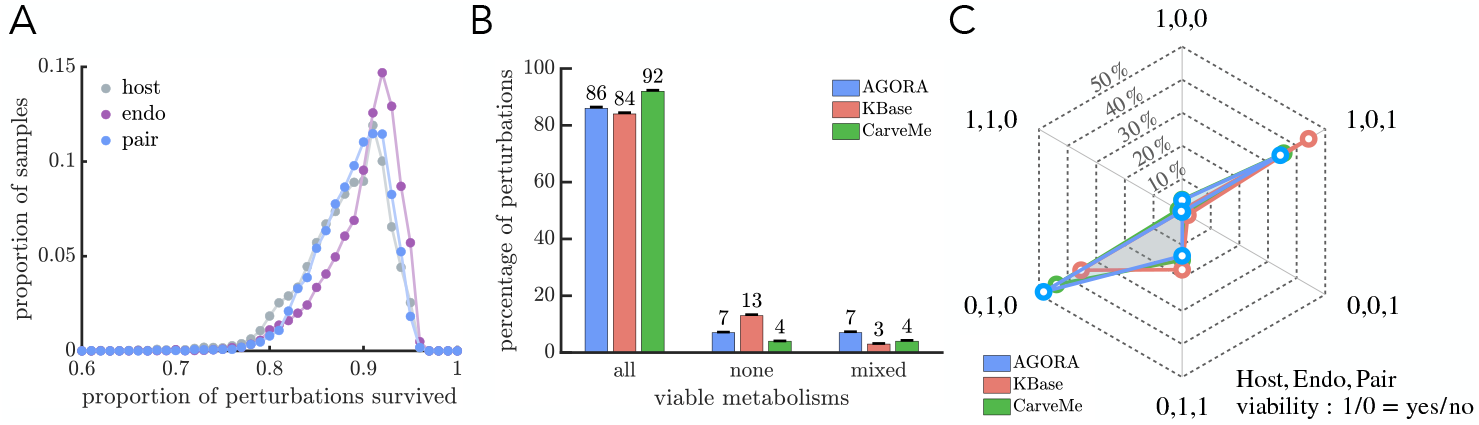
Survival competition between endosymbiosis and ancestral metabolisms in response to environmental degradation. A) The distributions show the robustness of 10,000 endosymbiosis metabolisms and their ancestral metabolisms sampled from the AGORA database (see Figure S3 for other databases). Robustness is quantified as the proportion of environmental perturbations survived by a metabolism. The distributions have similar single peaked shapes with *>* 80% of the population between the ranges of 85 − 95%. B) The bar graphs show the percentage of environmental perturbations for which all three metabolisms are viable, nonviable, or mixed across the three databases. In only 3 − 7% of perturbations is there a difference in the viability of the endosymbiosis metabolism compared to at least one of its ancestral metabolisms. C) The star plot displays the relative frequency of the possible outcomes for the mixed cases from B. Of the 6 possible scenarios for mixed viability, the two most frequent feature different survival between the ancestral endosymbiont metabolism compared to the other two, though which is more robust depends on the database (see Figure S4). The star plot also shows that the endosymbiosis survives more environmental perturbations than the ancestral host metabolism in all databases.

**Figure 5:**
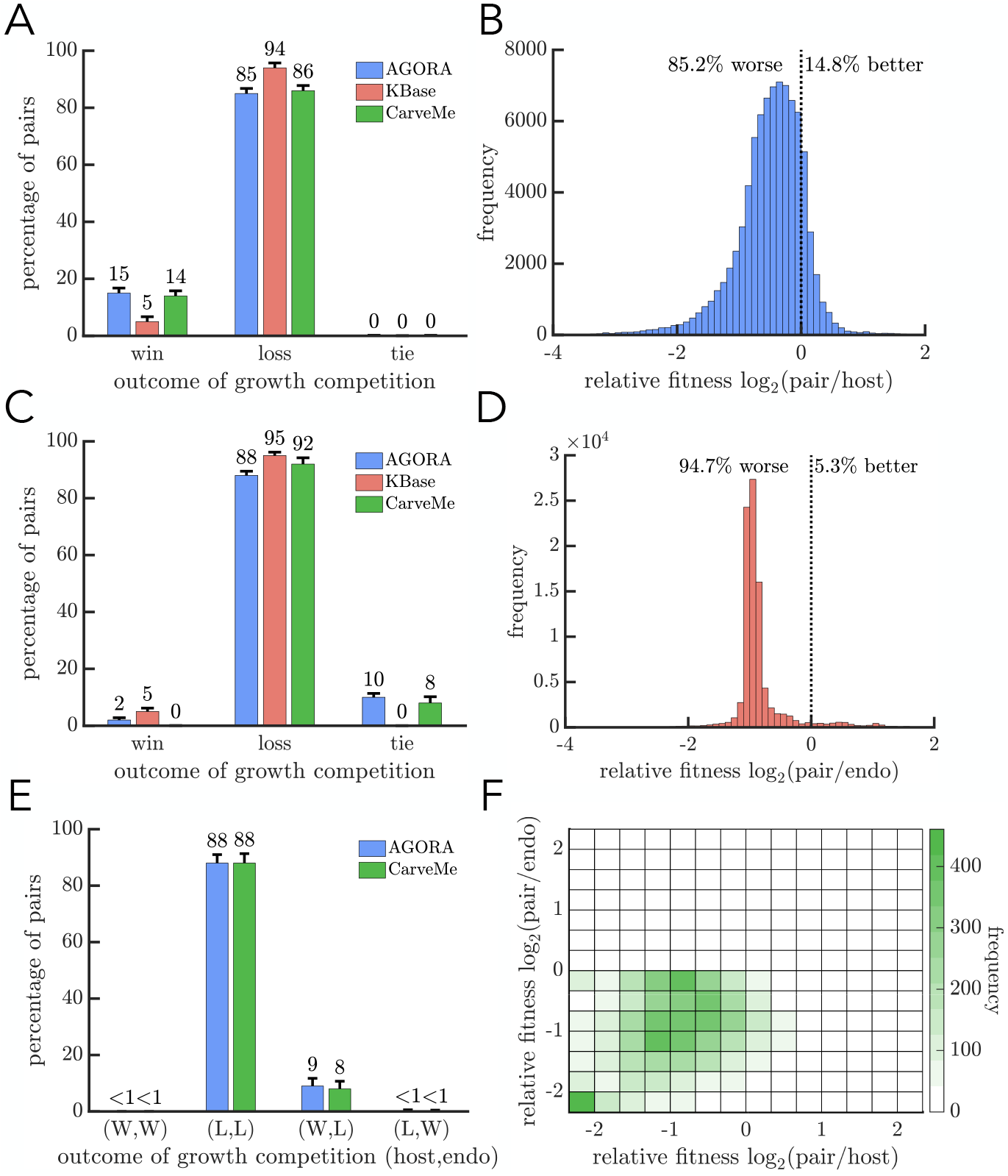
Growth-rate competition between endosymbioses and ancestral metabolisms. A) The bar graph shows the result of a growth-rate competition between the endosymbiosis and the ancestral host metabolism. The host grows faster for a majority of comparisons (85 − 94%). B) The histogram shows the relative growth rates of endosymbioses versus their ancestral host metabolisms, sampled from the AGORA database (see Figure S6 for other databases). When the endosymbiosis grows faster than its ancestral host cell, the fitness advantage is typically smaller in magnitude compared to its fitness disadvantage when it grows more slowly. C) The bar graph is similar to A but the comparison is between the endosymbiosis and the ancestral endosymbiont metabolism. Again the endosymbiosis grows more slowly in the majority of comparisons (88−92%). D) The histogram shows the relative growth rates of endosymbioses versus their ancestral endosymbiont metabolisms, sampled from the KBase database (see Figure S7). As in B, the typical growth advantage is smaller in magnitude than the growth disadvantage for endosymbioses. E) The bar graph shows the result of a growth-rate competition between the endosymbiosis and its ancestral metabolisms when all share the same environment. In both databases the most likely scenario is that the endosymbiosis grows more slowly than both ancestral metabolisms (88%). F) The relative fitness of the endosymbiosis versus its ancestral metabolisms from E is plotted using the CarveMe database (see Figure S8 for the AGORA database).

Since an endosymbiosis has to sustain two metabolisms, it could be that comparing its growth rate to those of an isolated ancestral host or endosymbiont metabolism is not a fair comparison. We address this possible issue by computing the growth rates of all three metabolisms when grown together in the same environment (see Methods: Growth rate calculation of communities). In short, we maximize each metabolism’s growth rate assuming that the total flux of metabolites through the system is maximal. We restrict our analyses to models from AGORA and CarveMe because models from the KBase database often yield multiple possible growth rates. Figure 5E shows that in the majority of cases (88% of samples) the host-endosymbiont pair grows more slowly than both ancestral metabolisms. It is rare (< 1%) that the host-endosymbiont pair can grow faster than the ancestral endosymbiont metabolism. In the 8 − 9% of samples where the host-endosymbiont pair grows faster than the ancestral host metabolism, the growth rate advantage is smaller in magnitude than its concomitant disadvantage compared to the ancestral endosymbiont metabolism (see Figure 5F).

Even if endosymbioses grow more slowly than their ancestors, they may be competitive if they are more adaptable. We evaluate this possibility by determining the effects of mutations on metabolisms when they are independent or in an endosymbiosis (see Methods: Adaptability assessment). Since we are interested in adaptability we focus on mutations that increase the bounds on reaction fluxes, which will either have no effect on a metabolism’s growth rate (neutral mutations) or increase it (beneficial mutations). We randomly sample 1000 endosymbioses that grow more slowly than their ancestors and compute the effects of all possible single mutations in host and endosymbiont reactions. We identified beneficial mutations in models from the AGORA and CarveMe databases but found none in models from KBase, because reaction bounds do not limit growth in KBase models. The adaptability analyses will thus rely only on models from the AGORA and CarveMe databases, though we did find similar results in KBase after imposing reaction bounds that do limit growth (see Methods: Adaptability assessment).

For mutations in host reactions, we find a similar number are beneficial in an endosymbiosis as in the ancestral host metabolism (Wilcoxon signed rank test, p≈ .07 in CarveMe and p≈ .27 in AGORA, see Figure 6A). In contrast, mutations in endosymbiont reactions are less often beneficial in an endosymbiosis than in the ancestral endosymbiont metabolisms (Wilcoxon signed rank test, p< 10^−6^ for both databases, see Figure 6B). Of the mutations that are beneficial in an endosymbiosis, few increase the growth rate beyond those of the ancestral metabolisms—only 5.0 − 5.6% of beneficial mutations in host reactions and 17.0 − 18.1% in endosymbiont reactions. Moreover, the majority of endosymbioses do not have any mutations that increase their growth rate above their ancestral metabolisms (see Figure 6C & D). It could be possible that, though rare, an endosymbiosis may have access to mutations that offer substantial growth benefits compared to its ancestral metabolisms. We evaluated this possibility by comparing the maximal growth rate reached by mutations in the ancestral metabolisms versus the endosymbiosis and found no such example in either database.

**Figure 6:**
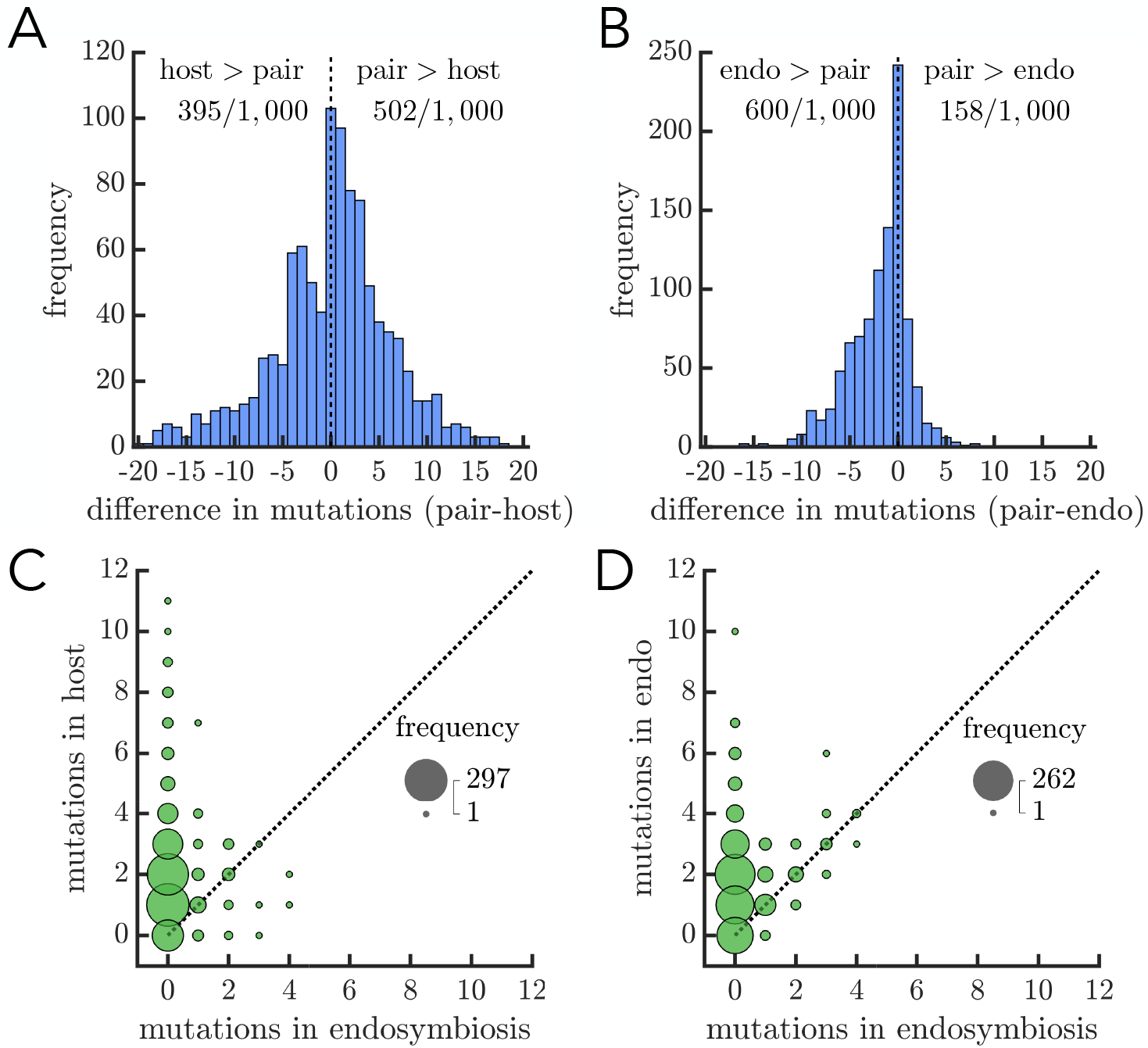
Adaptability of endosymbioses versus ancestral metabolisms. A) The histogram shows the difference in the number of beneficial mutations between the endosymbiosis and its ancestral host metabolism (data from AGORA database, see Figures S9 and S10 for CarveMe data). All mutations occur in host metabolism reactions and are deemed beneficial if they increase the growth rate. A sign-rank test supports the null hypothesis that the endosymbiosis and its ancestral host metabolism do not systemically differ in the number of beneficial mutations. B) The histogram is similar to A but compares the endosymbiosis and its ancestral endosymbiont metabolism for mutations in their shared reactions. Here, a sign-rank test rejects the null hypothesis, such that the ancestral endosymbiont metabolism has more beneficial mutations (*p <* .01). C) A bubble chart shows the frequency of mutations, in endosymbiosis vs ancestral host metabolisms, that increase the growth rate above that of ancestral host metabolism (data from CarveMe, see Figures S11 and S12 for AGORA data). There are more such mutations in ancestral host metabolisms than those in endosymbioses. D) The plot is similar to C except it is in terms of the ancestral endosymbiont metabolism. Again, there are more such mutations in ancestral endosymbiont metabolisms than those in endosymbioses.

## Discussion

An endosymbiosis between two prokaryotes gave rise to eukaryotes and likely facilitated the evolution of large complex organisms, but this is one of only a few examples of prokaryotic endosymbioses. While studies have identified possible obstacles to the formation of prokaryotic endosymbioses, there is a lack of quantitative frameworks that allow us to estimate their influence. Here we introduce a quantitative framework using genome-scale metabolic networks that evaluates the role of metabolic compatibility in the different stages of prokaryotic endosymbiosis evolution: viability, persistence, and evolvability. We find that over half of random pairs of metabolisms can form viable endosymbioses; however, the resulting endosymbioses often face fitness costs in terms of growth that they are unlikely to overcome through mutations.

A major result of our work here is that in the first stage towards a successful endosymbiosis— initial viability— metabolic compatibility produced only a very limited barrier, with over half of endosymbiotic configurations being viable. On the one hand, this is surprising given the substantial biochemical and metabolic diversity of prokaryotes. On the other hand, all cells share a metabolic core (e.g. the TCA cycle), facilitating the compatibility of metabolic networks [35, 36, 37]. Additionally, syntrophies and mutualisms in prokaryotes are commonplace, highlighting the relatively high likelihood for metabolic compatibility between prokaryotic cells configured in extracellular arrangements [38]. Examples include the occurrence of remarkably sophisticated *exo*-symbioses, such as aggregates of archaea and bacteria in which anaerobic methane oxidation is apparently achieved by passing an electron extracellularly [39, 40], close association symbioses in biofilms and microbial mats (e.g. [41, 42]), and ecosystems defined by effective supermetabolisms (e.g. [43]). Similarly, the frequent exchange of entire metabolic pathways via horizontal gene transfer [44, 45, 46] and the widespread success of plasmids in bacteria [47], which can be thought of as genomic endosymbionts, also point to the frequency of successful metabolic pairings. Since over 84 percent of species pairs had at least one viable configuration, the small barrier to metabolic viability seems to reflect more the spatial structure of paired metabolic networks, rather than the composition of the networks per se.

While metabolic compatibility did not provide a significant challenge to viability, it did emerge as a substantial filter during the persistence and adaptability stages, when endosymbioses compete with their ancestral metabolisms. Whether metabolisms competed to grow the fastest or survive environmental perturbations, an endosymbiosis rarely held a competitive advantage over both of its ancestors (< 1% of cases). In addition, mutations in endosymbioses rarely produced a growth advantage over ancestral endosymbionts (17 − 18.1%, Fig. 6D) or ancestral hosts (5 − 5.6%, Fig. 6C) These outcomes may be surprising given that all-else-being-equal increased network size should confer increased robustness to environmental perturbations, increased growth rates, and increased capacity to adapt [4, 5, 48]. However, unlike their ancestral metabolisms, endosymbioses must satisfy the biomass requirements of two metabolisms. Within this context, it is then somewhat expected that an early endosymbiosis may be at a fitness disadvantage compared to its ancestors, especially in the absence of any mechanism to coordinate or divide labor. Furthermore, the need to satisfy two sets of biomass requirements can limit or mask the effects of beneficial mutations, making it difficult to overcome initial fitness disadvantages.

Examining the rare cases when an endosymbiosis has higher fitness than its ancestors can shed light on the conditions in which it can persist. When subjected to environmental perturbations, the endosymbiosis has a competitive advantage against its ancestral host species. Similarly, when a premium is put on fast growth, the endosymbiosis is more likely to outcompete its ancestral host species than the ancestral endosymbiont species. Taken together, these results suggest that if an endosymbiosis were able to persist it would likely replace its ancestral host. This result is supported by the current ecological distributions of microbial eukaryotes and prokaryotes. Microbial eukaryotes often coexist with the ancestral lineage of their mitochondrial endosymbiont, the alpha-proteobacteria [49, 50], but they seldom coexist with the ancestral lineage of the host, presumably of archaeal origin— likely a Lokiarchaeota [51, 52, 53, 54]. Moreover, our competition analyses suggest that while an endosymbiosis almost never grows faster than both of its ancestors, an appropriate combination of perturbed environments may allow a nascent endosymbiosis to survive when both its ancestors do not. These results are consistent with the argument that, compared to bacteria, eukaryotes may have been K-selected with slower rates of reproduction and limiting nutrient supplies [55]. The results are also corroborated by the observation that the few other documented prokaryotic endosymbioses are found in relatively nutrient-poor environments [56, 14, 15], e.g. the bristles of a scale worm near a deep hydrothermal vent [56] or in a specialized organ of a sap-feeding mealybug [15].

We have used our metabolic framework to make general predictions about the role of metabolic compatibility in shaping nascent prokaryotic endosymbioses. The quality of more specific predictions about particular species and environmental conditions will likely depend on the accuracy of metabolic network models. Here we drew conclusions using three different metabolic network databases; and although they broadly agreed about many aspects of metabolic compatibility in endosymbioses, there were some key differences. For example, the outcome of survival competition between a host-endosymbiont pair and its ancestral endosymbiont metabolism depended on the database: the endosymbiont won more often in the CarveMe and AGORA databases but not in KBase. Differences between databases could stem from many factors including the organisms, environments, or procedures used to construct the networks themselves. Indeed, research into metabolic network construction and validation is very active and there will likely be much future development. While we expect this future development to lead to refinements in our quantitative predictions, it is unlikely to change the broader conclusions that were common to the three databases considered here. Given the diverse formats and modes of construction of metabolic networks in the three databases, the common findings are likely to be universal features of metabolism. Regardless of this expectation, our metabolic endosymbiosis framework could readily accommodate future advances in the field of metabolic networks—it simply requires rerunning the computations with updated networks or different environmental conditions. Thus, our quantitative framework is adaptable and should continue to make useful predictions concerning the role of metabolic compatibility in the evolution and ecology of prokaryotic endosymbioses.

An overarching goal of our work has been to assess factors responsible for the relative rarity of prokaryotic endosymbioses. We have focused on metabolic compatibility because it was amenable to existing methodology and plays a role in the initiation, persistence, and adaptation of endosymbioses. While our results reveal significant limitations imposed by metabolic compatibility, it is unlikely that metabolic compatibility alone accounts for the relative rarity of prokaryotic endosymbioses. If it were the dominant barrier then it would imply that metabolic compatibility is far less often an issue when the host is a eukaryote. We can gain some insight into this possibility by considering the abundant examples of endosymbioses with eukaryote hosts [57, 58]. We expect that multicellular eukaryotes can more easily accommodate intercellular endosymbionts because they can manipulate the space between their cells to create specialized metabolic environments for their endosymbionts, such as light organs in bobtail squid [59, 60] or hindguts in termites [61]. If, instead, we consider only intracellular endosymbionts which can be found in both multicellular and unicellular eukaryotes [57, 58, 11, 62, 63, 64], then many of these also involve specialized spaces, e.g. bacteriocytes [65] or vacuoles [10]. The fact that so many eukaryotic endosymbioses involve specialized compartments suggests some adaptive benefit to controlling the metabolic exchanges between species, i.e. they may not be innately synergistic—though it is unclear whether these spaces evolved prior to or in conjunction with the endosymbiosis. Moreover, we may expect that the increased presence of structured compartments could create potential issues with transport, which was the primary source of nonviability in the initiation stage of our prokaryotic endosymbioses. Thus, it seems likely that some other factor(s) works in conjunction with metabolic compatibility to make prokaryotic endosymbioses rare.

Of the other factors limiting the emergence of prokaryotic endosymbioses, the one that has often been invoked as substantial is the difficulty of internalizing a cell within another in the absence of phagocytosis [16]. Indeed, the ubiquity of phagocytosis in eukaryotes creates significantly more opportunities to initiate an endosymbiosis, e.g. amoeba frequently engulf different bacteria that can resist digestion [66]. However, phagocytosis may not be the only physiological mechanism for a cell to internalize another cell (e.g. [67, 16, 34]). Many other studies have demonstrated the plasticity of prokaryote membranes and cytoskeletal architecture [68, 69, 51, 70] suggesting the absence of phagocytosis may not be as dominant a barrier as once thought. And the continuing discovery of new functions, traits, and morphologies in prokaryotes also emphasize the importance of evaluating multiple barriers, not just phagocytosis. Indeed, remarkably, phylogenetic analysis of prokaryote endosymbiosis within mealybug cells suggests that the endobacterial symbiosis may have originated multiple times during mealy bug diversification [15], despite the prokaryote host lacking an obvious capacity for endocytosis. It thus seems highly probable that many more similar examples of such endobacteria symbionts remain to be discovered, especially given that this prokaryotic endosymbiosis was previously overlooked [71] and the microbiomes of millions of invertebrate species have not been investigated.

Ultimately, given the functional diversity and abundance of microbes on Earth, the expanding list of discoveries that soften phagocytosis as a barrier, and the metabolic viability of many potential endosymbiosis, a reasonable interpretation of our results is that hundreds to thousands or more prokaryote endosymbioses remain to be discovered. However, the reduced metabolic robustness, growth rates, and evolvability that these endosymbioses likely face severely limits their persistence and potential to radiate spectacularly. The high rates of extinction and low rates of speciation resulting from these metabolic considerations would sustain only a relatively low diversity of extant prokaryotic endosymbioses confined to environments in which they are less likely to be outcompeted by ancestral host or endosymbiont lineages, such as nutrient-poor environments.

## Methods

### Genome-scale metabolic model curation

We obtained metabolic models of diverse prokaryotes in .xml format made available in [72] from the AGORA[73], KBase[74], and CarveMe[75] databases (see Supplementary Table S1 for details). We used the COnstraint-Based Reconstruction and Analysis Toolbox (COBRA) [76] to create .mat files for further analysis using MATLAB [77]. The .mat format of a metabolic model includes a stoichiometric matrix of compounds and reactions, a list of compound names, lower and upper bounds for reaction fluxes, an objective function that identifies the biomass reaction, a right hand side vector that indicates how each compound is balanced, and other identifying information. Metabolic models are partitioned across two compartments (cytoplasm [c] and extracellular [e]) except those in CarveMe which include an additional periplasm [p] compartment. All models come with an implicit environment—determined by the bounds on reaction fluxes and right hand side vectors—that make a set of extracellular compounds accessible so the metabolism is viable, i.e. it can synthesize all of its biomass compounds. Determining whether a metabolism is viable requires performing flux balance analysis and solving the associated linear program. We used the Gurobi optimization software [78] to solve all linear programs and confirmed that all metabolic models are initially viable, where viability is defined as having an objective function value (growth rate) above a tolerance of 0.001. For each database, we restructured the metabolic models to facilitate analyses of possible endosymbioses. First, we identified all compounds and reactions used within a database and reformatted the stoichiometric matrices (*S*) for the metabolic models so that the same row corresponds to the same compound across models. Second, we changed how environmental compounds are made available to metabolic models. Initially, metabolic models access environmental compounds through source/sink reactions whose upper and lower bounds on flux determine how much of the compound is available. We replaced these reactions with mathematically equivalent constraints on the amount of compound available. So a source/sink reaction flux *r*_*i*_ for compound *c*_*j*_ with lower and upper bounds *l*_*i*_ and *u*_*i*_ respectively would be replaced with the equivalent constraint on the compound concentration, *l*_*i*_ ≤ *c*_*j*_ ≤ *u*_*i*_, note *c*_*j*_ = Σ_*k*_ *S*_*j,k*_*r*_*k*_. This modification makes it easier to create joint environments using different metabolic models. Finally, for each metabolic model, we created different stoichiometric matrices depending on its role as either host or endosymbiont. When the metabolism is a host, its stoichiometric matrix (*S*_*H*_) is the same as if it were growing in isolation. When the metabolism is an endosymbiont, its extracellular compartment is identical to the cellular compartment of the host. Thus, we partitioned the stoichiometric matrix depending on whether the compounds are strictly within the endosymbiont (*S*_*E*_) or within the host (*S*_*E*→*H*_).

### Assessing growth and viability of metabolisms

We compute the growth rate of metabolisms, either in isolation or in endosymbioses, by performing flux balance analysis and solving the associated linear program. If the index for the biomass reaction is *λ* then the linear program for its growth in isolation is:

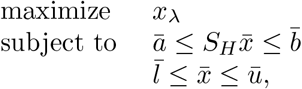

where 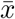 is a vector of reaction fluxes, 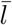 is a vector of lower bounds for fluxes, *ū* is a vector of upper bounds for fluxes, *S*_*H*_ is the stoichiometric matrix, *ā* is a vector of lower bounds for compound concentrations, and 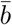 is a vector of upper bounds for compound concentrations.

For an endosymbiosis, we denote the fluxes of the host as 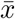 and those of the endosymbiont as 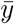. We use the *H* and *E* indices to indicate host or endosymbiont respectively. The linear program for two metabolisms in an endosymbiosis is then:

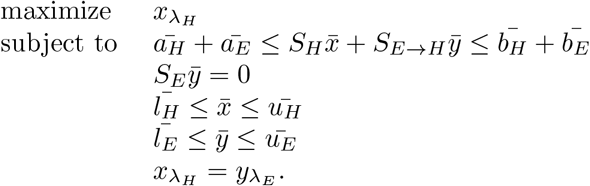

The 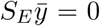 condition stems from the fact that compounds in the cytoplasm and periplasm compartments are balanced in the endosymbiont. The last condition 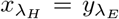 requires that the host and endosymbiont grow at the same rate.

### Fixing viability

We attempted to fix nonviable endosymbioses through a multi-step process. First, we determined whether the endosymbiosis could be viable if all transport between the cytoplasm and extracellular compartments are available. We identified all environmental compounds that 1) are available in the environment with nonzero amounts, 2) exist in both environmental and cellular compartments and 3) are usable by the endosymbiont such that at least one reaction has it. We then added extra reactions in the host-endosymbiont joint stoichiometric matrix that enabled those compounds to be transported between the host’s [e] and [c] compartments. We performed flux balance analysis to determine the growth rate. If the growth rate was below the tolerance .001 it meant that the lack of viability was also due to the endosymbiont not having access to the external environment. We then confirmed that viability could be restored by providing the endosymbiont access to the external environment through creation of source/sink reactions in *S*_*E*_ for compounds present in the environment at nonzero amounts. All nonviable endosymbioses could be fixed by a combination of restoring transport and allowing the endosymbiont to access extracellular compounds. In cases where viability could be restored by providing transport reactions alone, we estimated a potential minimal set of compounds following the algorithm outlined in Figure S5.

### Growth rate calculation of communities

We compute the growth rates of the host-endosymbiont pair together with its ancestral metabolisms in the same environment through a two-step process. First we compute the maximal flux through the entire community by maximizing the sum of fluxes through the three biomass reactions: ancestral host 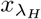, ancestral endosymbiont 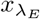, and host-endosymbiont pair 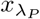. Second we maximize the flux through each of the three biomass reactions one at a time while requiring the community flux be held constant at its maximal value (call it *z*), i.e. we add the constraint 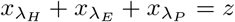. We perform the second step because the solution to a linear program may not be unique, so there may be multiple partitions of growth within a community that give rise to the same total community flux. For our analyses we considered only those cases in which the growth rates of the host-endosymbiont pair and its ancestral metabolisms are unique when the total community flux is maximal. We determined uniqueness by requiring that the computed growth rates for metabolisms did not vary above 10% between its highest and lowest values. We found that the AGORA and CarveMe databases frequently gave unique values, 99.7% and 98.2% of computations respectively; however, KBase only met this criteria for uniqueness in 0.02% of computations. Despite the lack of uniqueness in the KBase computed growth rates, the maximal host-endosymbiont pair growth rate rarely exceeded the growth rates of both its ancestral metabolisms, 3 out of 10000 computations.

### Adaptability assessment

We evaluate the adaptive potential of metabolisms by determining how the computed growth rate is affected by mutations that increase the bounds of reactions. Since increasing the bounds of reactions can never decrease growth rates, the mutations are either neutral or beneficial. For ancestral metabolisms, either host or endosymbiont, we systematically modify each reaction bounds one at a time by a factor of 1000. Thus if a reaction flux *x*_*i*_ has a lower bounds *l*_*i*_ (where *l*_*i*_ ≤ 0) and an upper bound *u*_*i*_ then we transform it the bounds to 1000*l*_*i*_ ≤ *x*_*i*_ ≤ 1000*u*_*i*_. We compute the growth rate following this mutation and then reset the bounds to evaluate another mutation. For host-endosymbiont pairs we evaluate both mutations in endosymbiont reactions and host reactions separately.

While our methodology to assess adaptability was able to identify mutations that affected growth rate in the AGORA and CarveMe databases, it did not find any in KBase. The lack of any beneficial mutations in KBase suggests that the reaction bounds do not constrain the growth rate. We tested this hypothesis by decreasing the bounds of all reaction rates by a factor of 100 and re-computing growth rates. We found that that the growth rates did decrease and we could identify beneficial mutations in the metabolisms of the host-endosymbiont pair as well as its ancestors. The results are similar to the analyses with AGORA and CarveMe described in the main text: mutations were more often beneficial in ancestral metabolisms than host-endosymbiont pairs (87.7% for host reactions and 78.0% for endosymbiont reactions).

## Supplementary Material

### Database information

**Table S1:** Summary data for metabolic model databases. AGORA contains metabolic reconstructions of human gut bacteria comprising over two hundred genera [73]. KBase is a platform created by The United States Department of Energy for sharing, integrating, and analyzing data of communities of plants and microbes [74]. CarveMe is a tool to automate the construction of metabolic models and circumvent issues with manual curation [75]. Due to its automated nature, it can used to generate large collections of metabolic models. Together these three databases represent a large and varied collection of metabolic models.

| Database | Number of metabolisms | Number of compounds | Number of reactions |
| --- | --- | --- | --- |
| AGORA | 818 | 2072 | 3815 |
| KBase | 1637 | 1675 | 3245 |
| CarveMe | 5587 | 2340 | 4245 |

### Predicting viability

In this section, we assess whether simple heuristics concerning the similarity of host and endosymbiont metabolic networks may predict viability. We first turn to the biomass reactions which effectively list all of the compounds required by the host and endosymbiont in order to grow. If the biomass reactions contain completely different compounds then we might expect a potential endosymbiosis to be nonviable because the host and endosymbiont have different metabolic needs. For example, if the endosymbiont requires a compound that the host does not need then the host may lack the ability to transport it from the external environment into its cytoplasm where the endosymbiont can access it. In contrast, if the biomass reactions contain the same compounds then both metabolisms need to make the same products, which would increase the likelihood that the host transports compounds needed by the endosymbiont. We predict then that the number of compounds from the endosymbiont’s biomass reaction that are missing in the host, may increase the likelihood of nonviability. Figure S1 shows that the AGORA database matches this expectation to some extent—though the relationship is not strictly monotonic. In the KBase database the probability of a viability is actually higher if one compound is missing rather than none. In the CarveMe database, all biomass reactions contain the same compounds so the metric of missing compounds is not useful in predicting nonviability (plot not shown).

**Figure S1:**
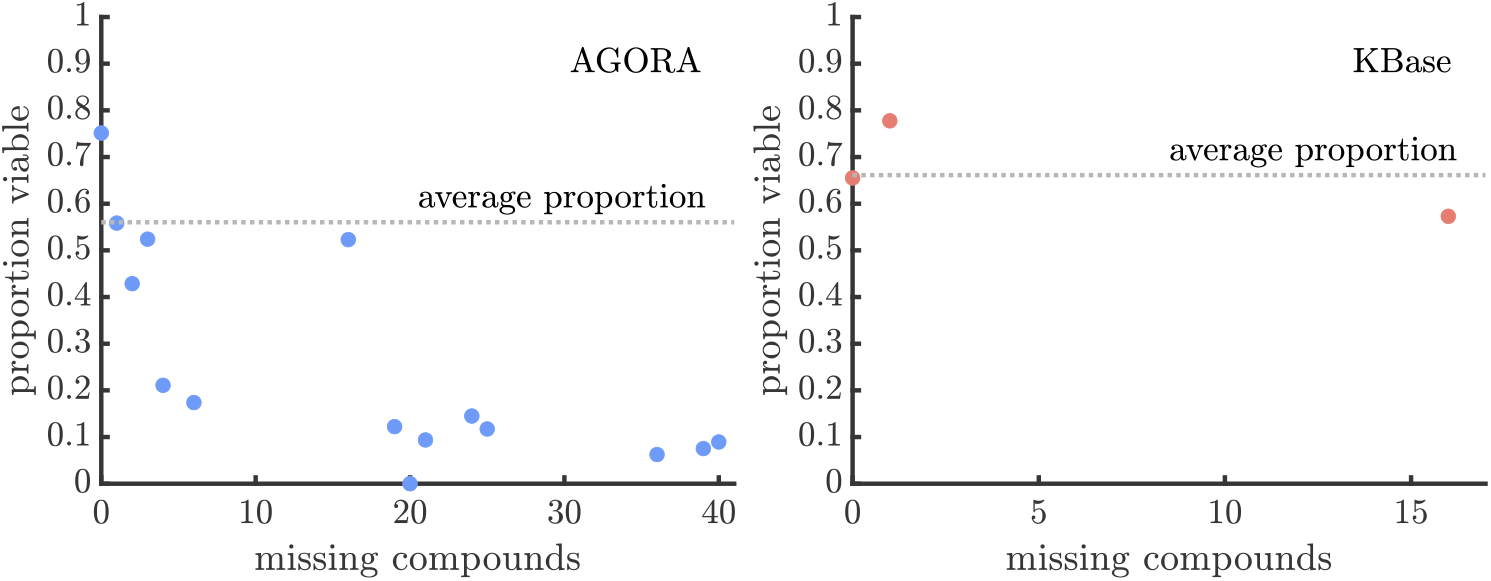
Proportion of missing biomass compounds as an indicator of nonviability. Each plots shows the proportion of endosymbioses that are viable as a function of the number of compounds in an endosymbiont’s biomass reaction that are missing from the host’s. The dashed line in each plot shows the average proportion of viable endosymbioses across all samples within a database. The AGORA plot suggests a relationship between the number of missing biomass compounds and nonviability. KBase biomass reactions belong to a few distinct classes which makes it difficult to discern any relationship. Indeed the pairs of metabolisms with one missing biomass compound are more viable than those with none. All CarveMe biomass reactions contain the same compounds so there are no missing compounds (plot not shown).

Next we consider whether the proportion of reactions shared between the host and endosymbiont is a predictor of viability. If the host and endosymbiont share many reactions then they may also share metabolic pathways and higher order metabolic structures, making them more likely to be metabolically compatible. Figure S2 shows the results of this analysis. The relationships vary across databases but once a host metabolism has ≥75% of an endosymbiont’s reactions there is an above average probability that they can form a viable endosymbiosis—although in KBase and CarveMe this relationship is not monontonic and even shows a dip when the host’s metabolism contains all of the endosymbiont’s reactions.

**Figure S2:**
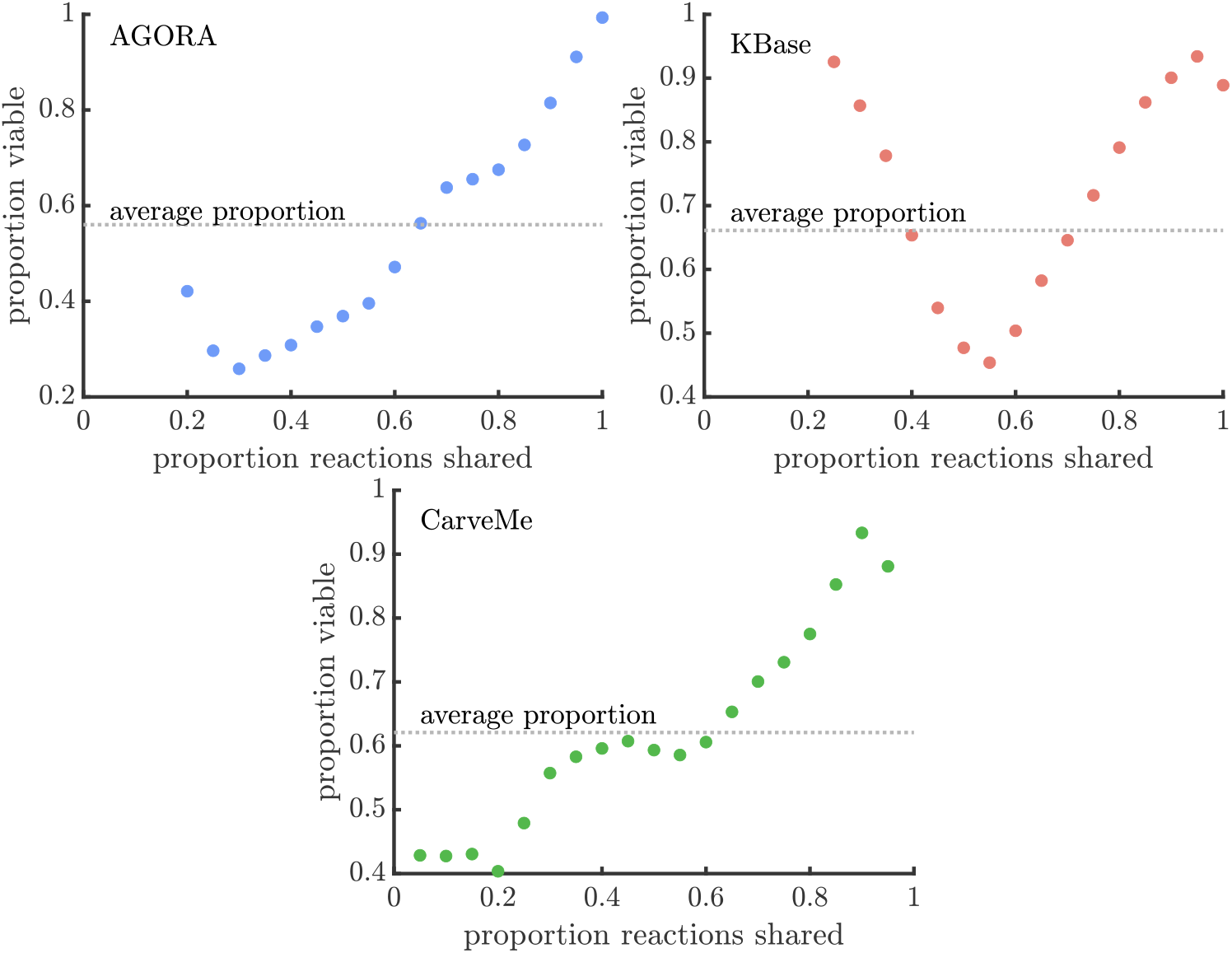
Proportion of shared reactions as an indicator of viability. Each plot shows the proportion of endosymbioses that are viable as a function of the proportion of an endosymbiont’s reactions that are shared with the host. The dashed line shows the average proportion of viable endosymbioses across all samples within a database. The plots show that the relationship between shared reactions and viability is not consistent across databases, e.g. KBase has a Ushaped relationship while CarveMe has a mostly monotonic relationship. One common feature is that if a host metabolism has ≥75% of the endosymbiont’s reactions then there is a higher than average probability that the resulting endosymbiosis is viable.

Finally, we note that if we use the same metabolic network for both host and endosymbiosis we still find nonviable pairings: 7/818 (or 0.86%) in AGORA, 211/1637 (or 12.89%) in KBase, and 459/5587 (or 8.22%) in CarveMe.

### Viability selection

**Figure S3:**
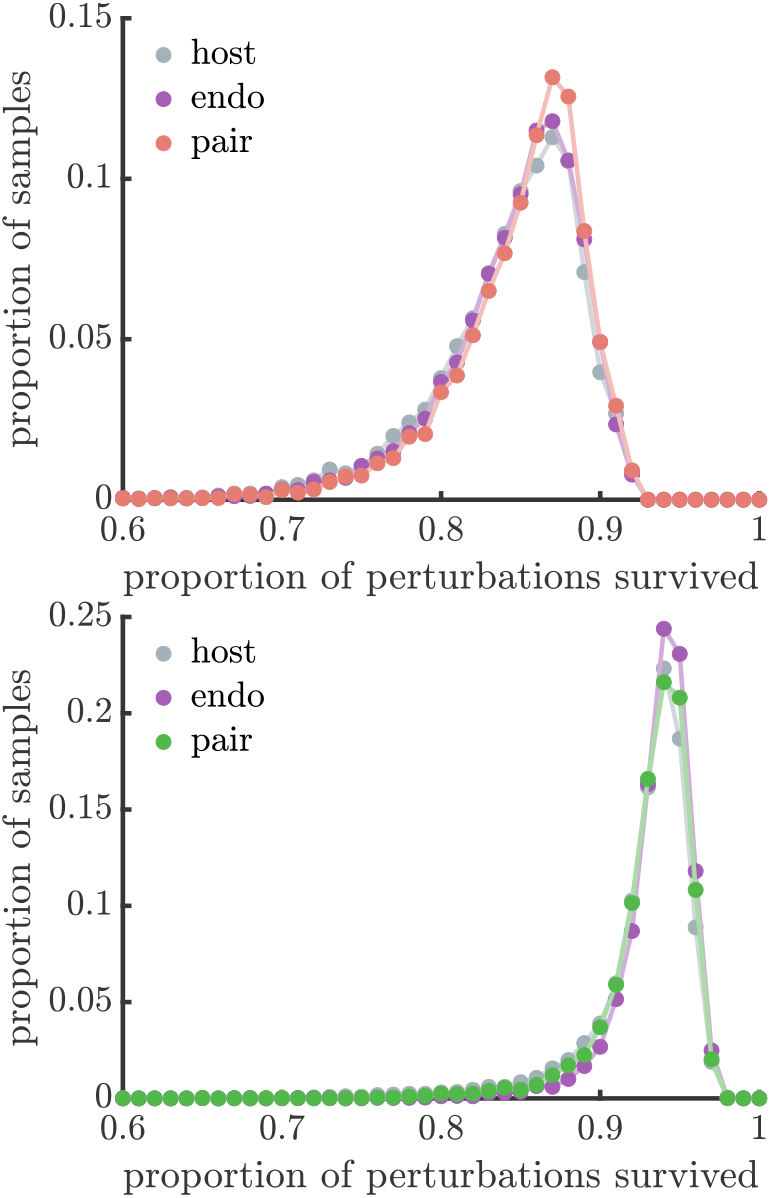
Companion to Figure 4A. The distributions show the robustness of 10,000 endosymbiosis metabolisms and their ancestral metabolisms sampled from the KBase database (top) and the CarveMe database (bottom). As in Figure 4A, robustness is quantified as the proportion of environmental perturbations survived by a metabolism. The distributions are similar in shape within databases.

**Figure S4:**
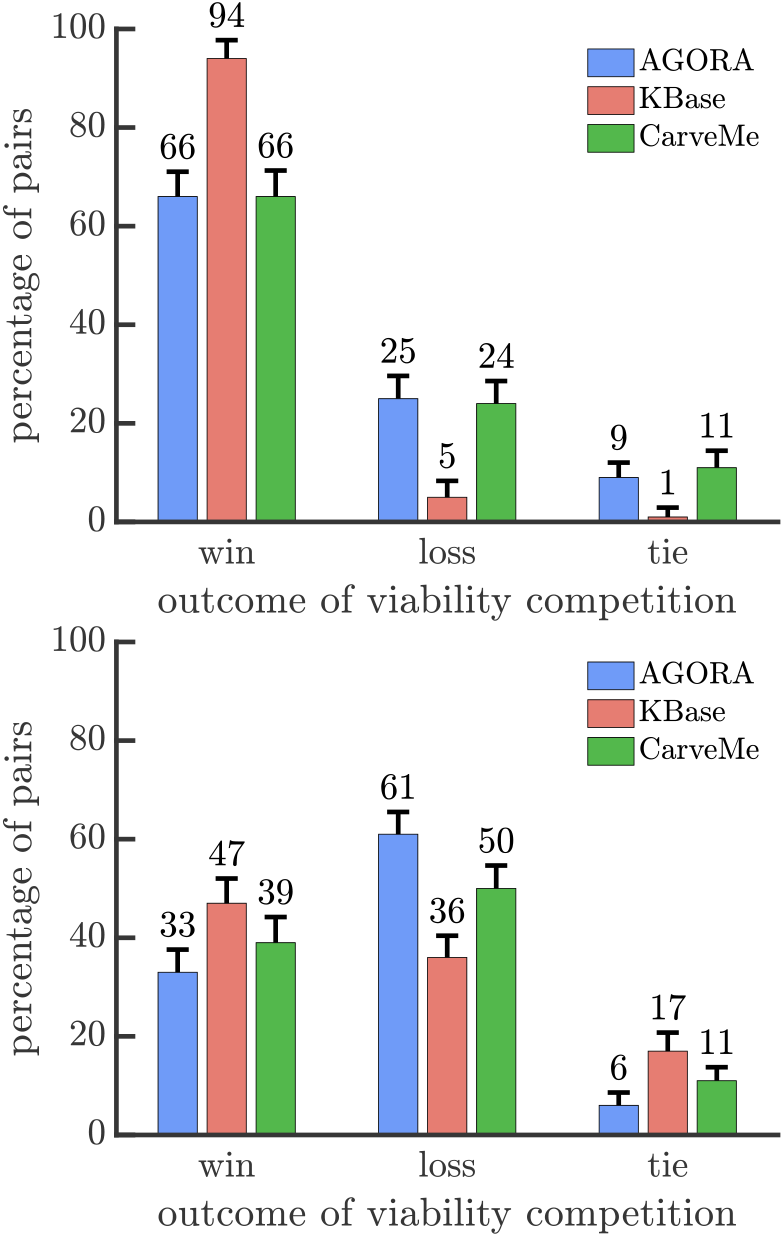
Companion to Figure 4C. (top) The bar graphs show the results of comparing the endosymbiosis with the ancestral host metabolism in terms of surviving environmental degradation. Across the databases the endosymbiosis is more robust to environmental degradation than the ancestral host metabolism. (bottom) Similar to the top plot except it compares the endosymbiosis to the ancestral endosymbiont metabolism. The endosymbiosis is more robust than the ancestral endosymbiont metabolism in the KBase database and less robust in the AGORA and CarveMe databases.

**Figure S5:**
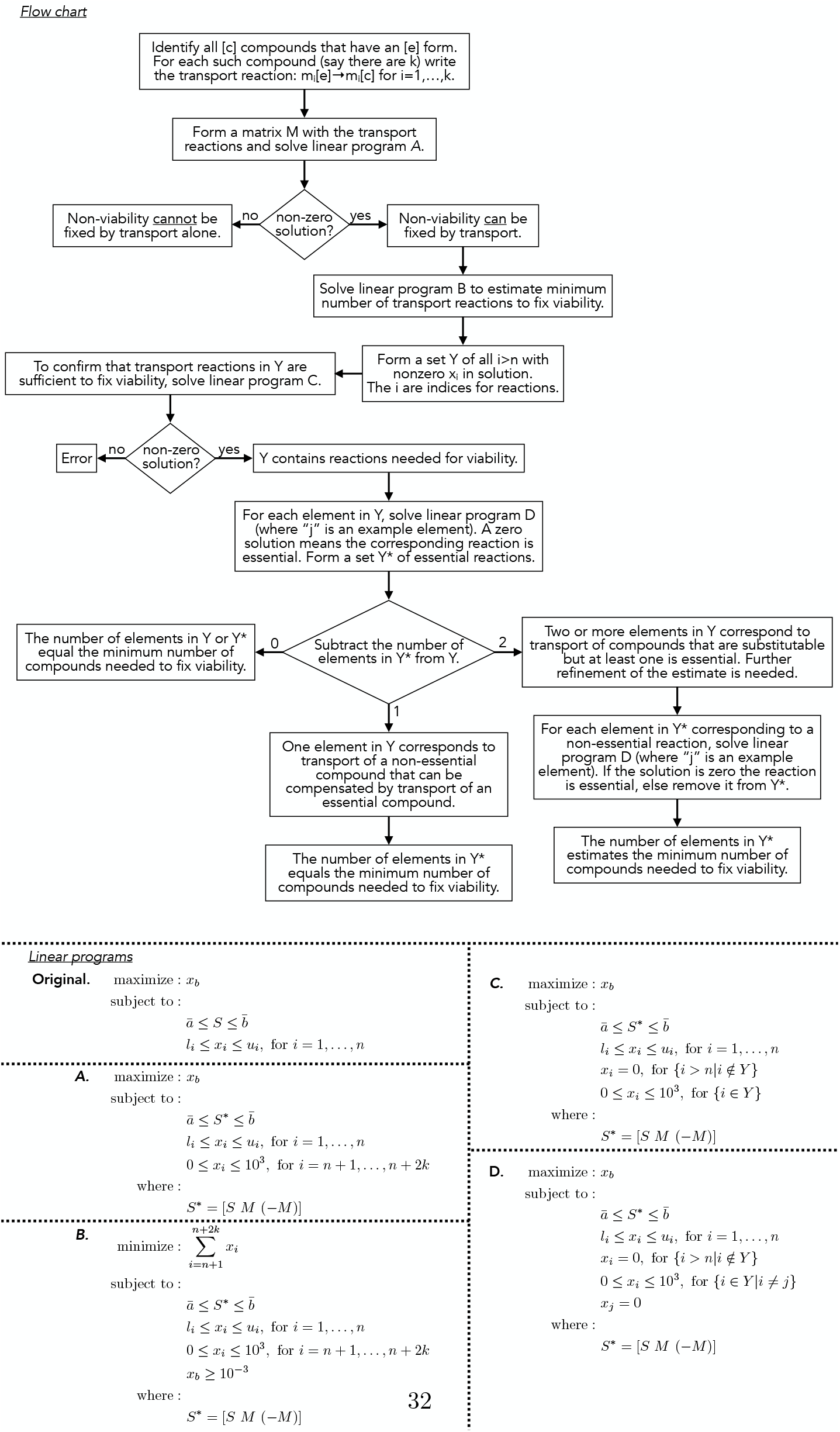
Flow chart for estimating minimum number of compounds needed to fix viability.

### Growth-rate selection

**Figure S6:**
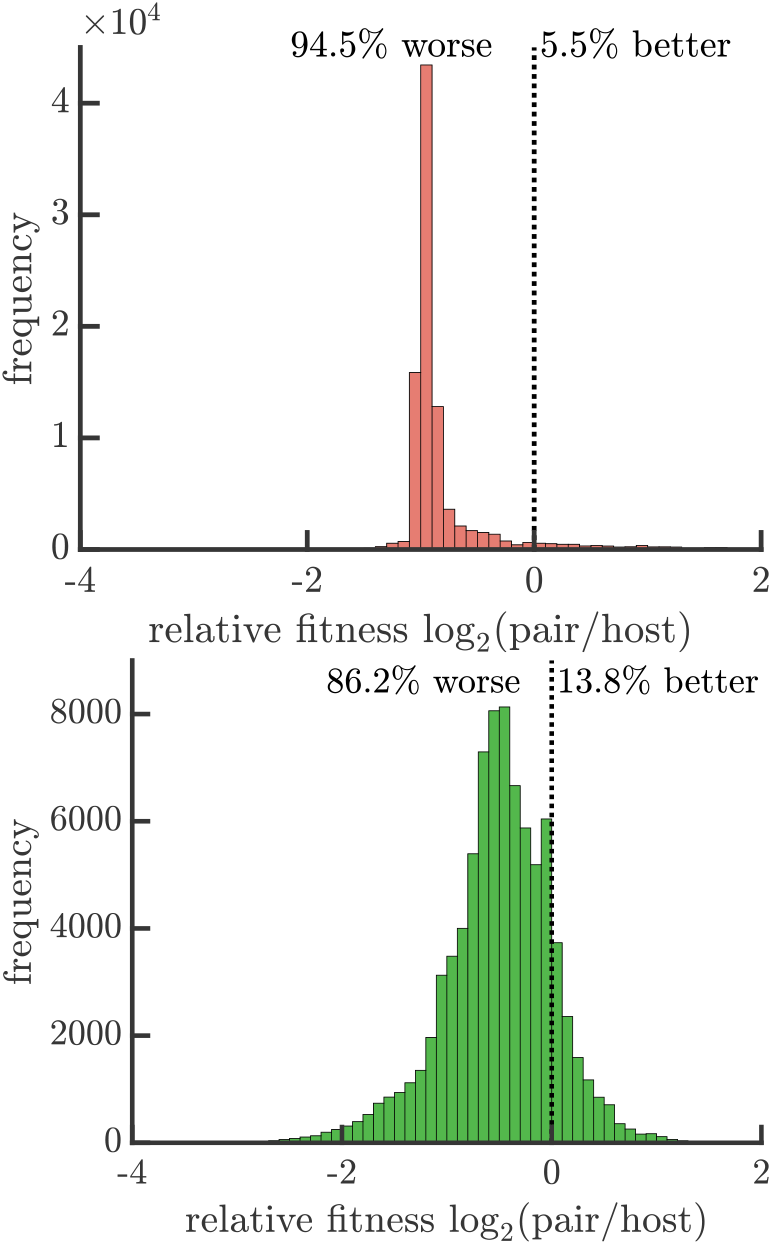
Companion to Figure 5B. The distributions are the same as in Figure 5B except instead of the AGORA database the data is from the KBase database (top) or the CarveMe database (bottom). Both show that the growth rate of host-endosymbiont pairs is more often less than the ancestral host metabolism.

**Figure S7:**
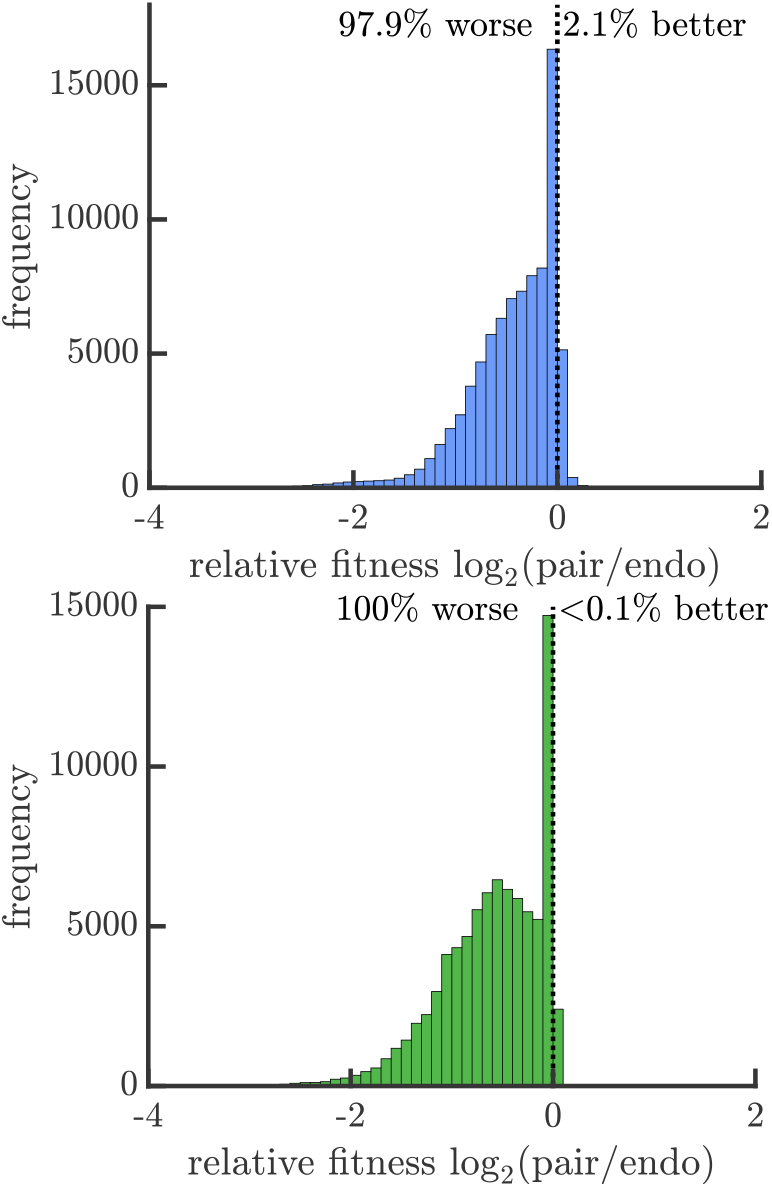
Companion to Figure 5D. The distributions are the same as in Figure 5D except instead of the KBase database the data is from the AGORA database (top) or the CarveMe database (bottom). Both show that the growth rate of host-endosymbiont pairs is more often less than the ancestral endosymbiont metabolism.

**Figure S8:**
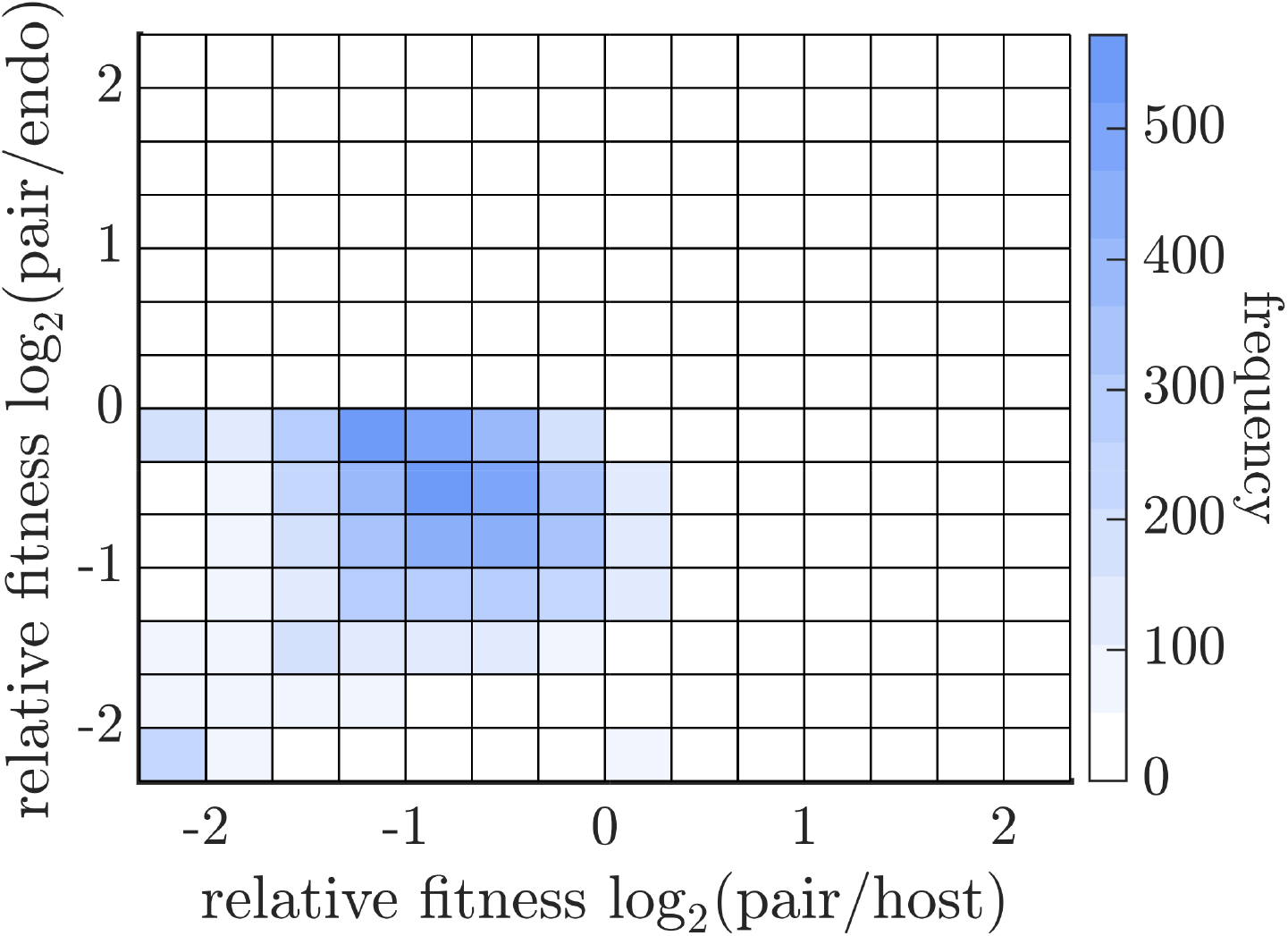
Companion to Figure 5F. The plot is the same as in Figure 4F except instead of the CarveMe database the data is from the AGORA database. We find a similar pattern in which the host-endosymbiont pair is less fit than both of its ancestral metabolisms.

### Adaptability figures

**Figure S9:**
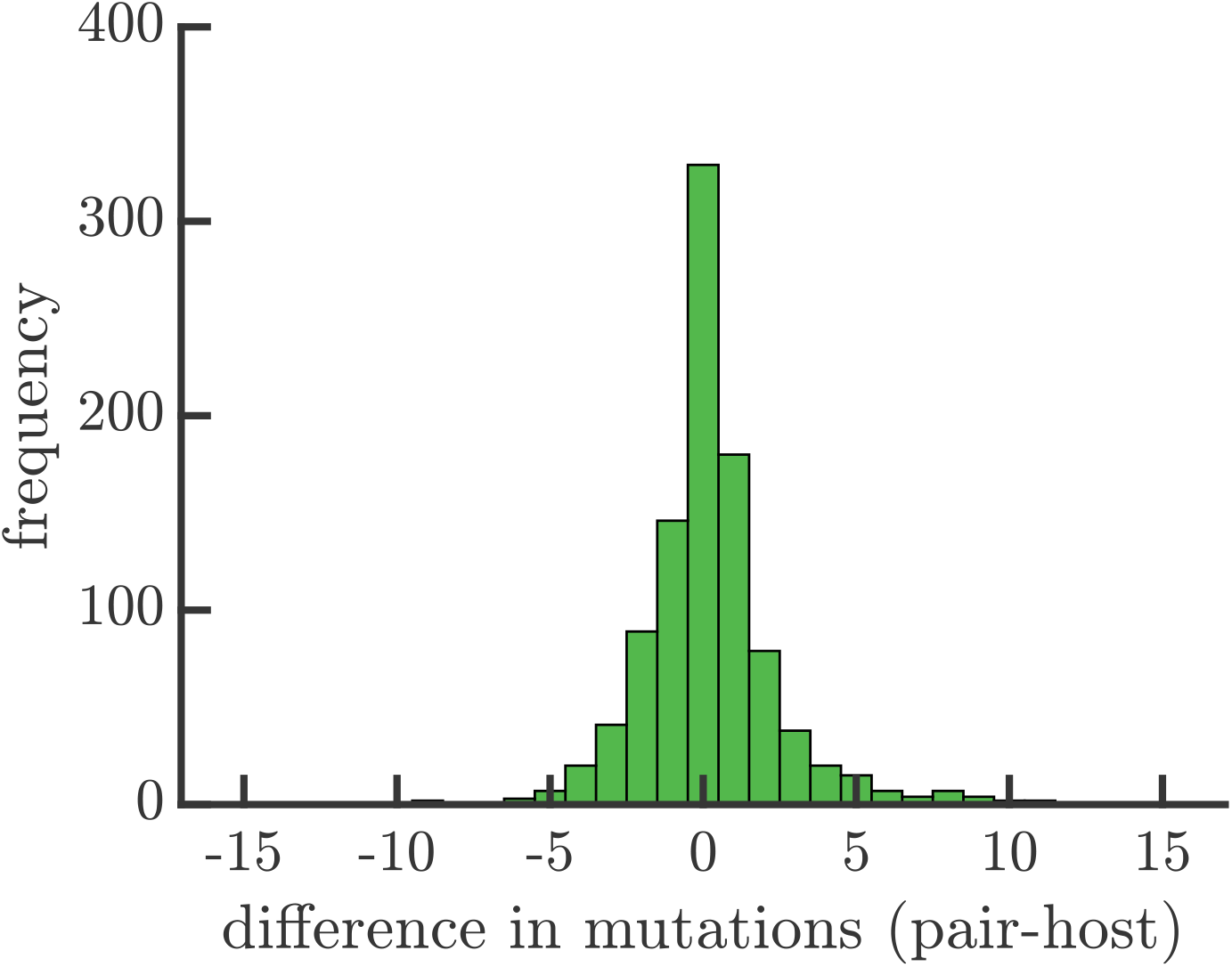
Companion to Figure 6A. The plot is the same as in Figure 6A except instead the data is from the CarveMe database instead of the AGORA database.

**Figure S10:**
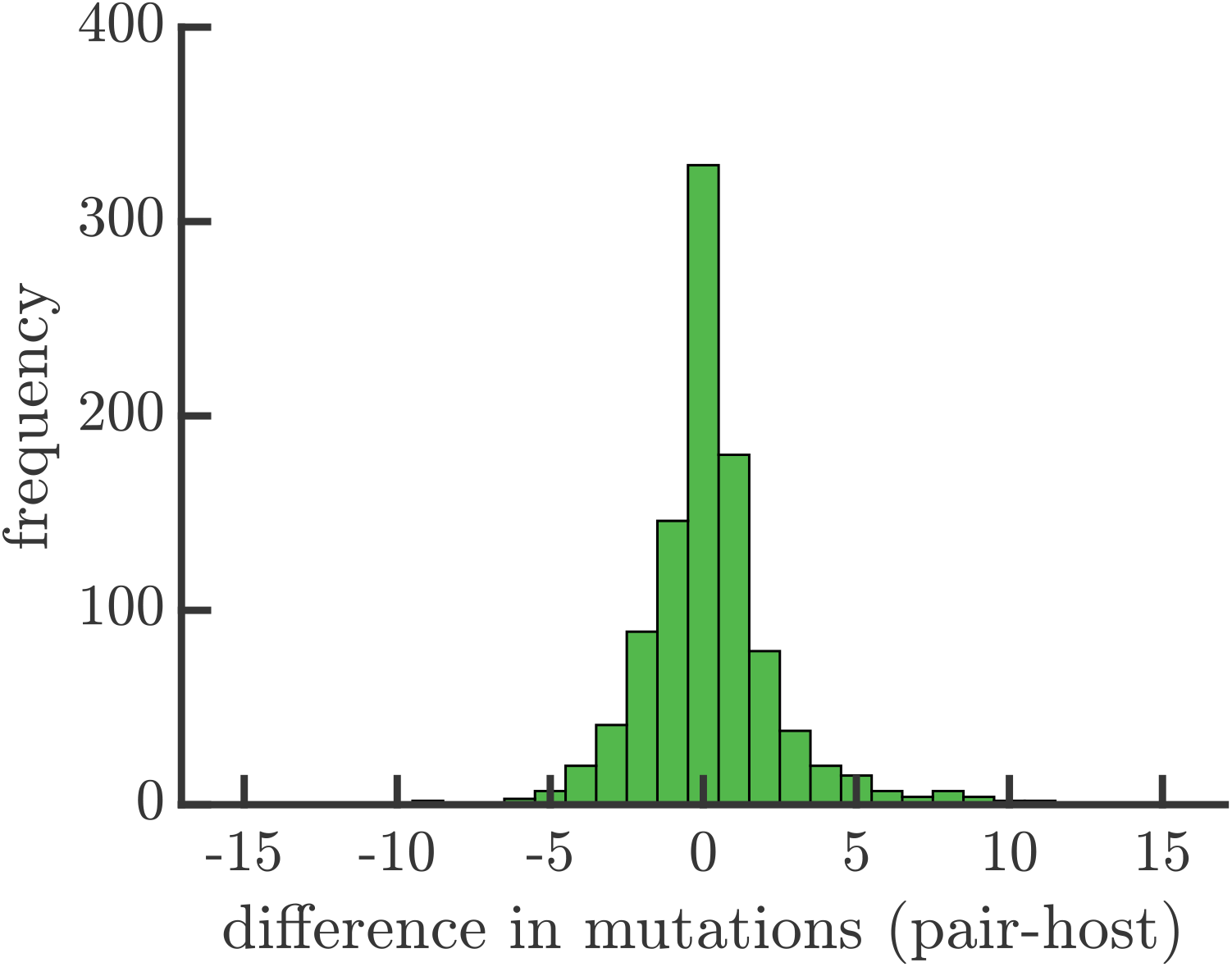
Companion to Figure 6B. The plot is the same as in Figure 6B except instead the data is from the CarveMe database instead of the AGORA database.

**Figure S11:**
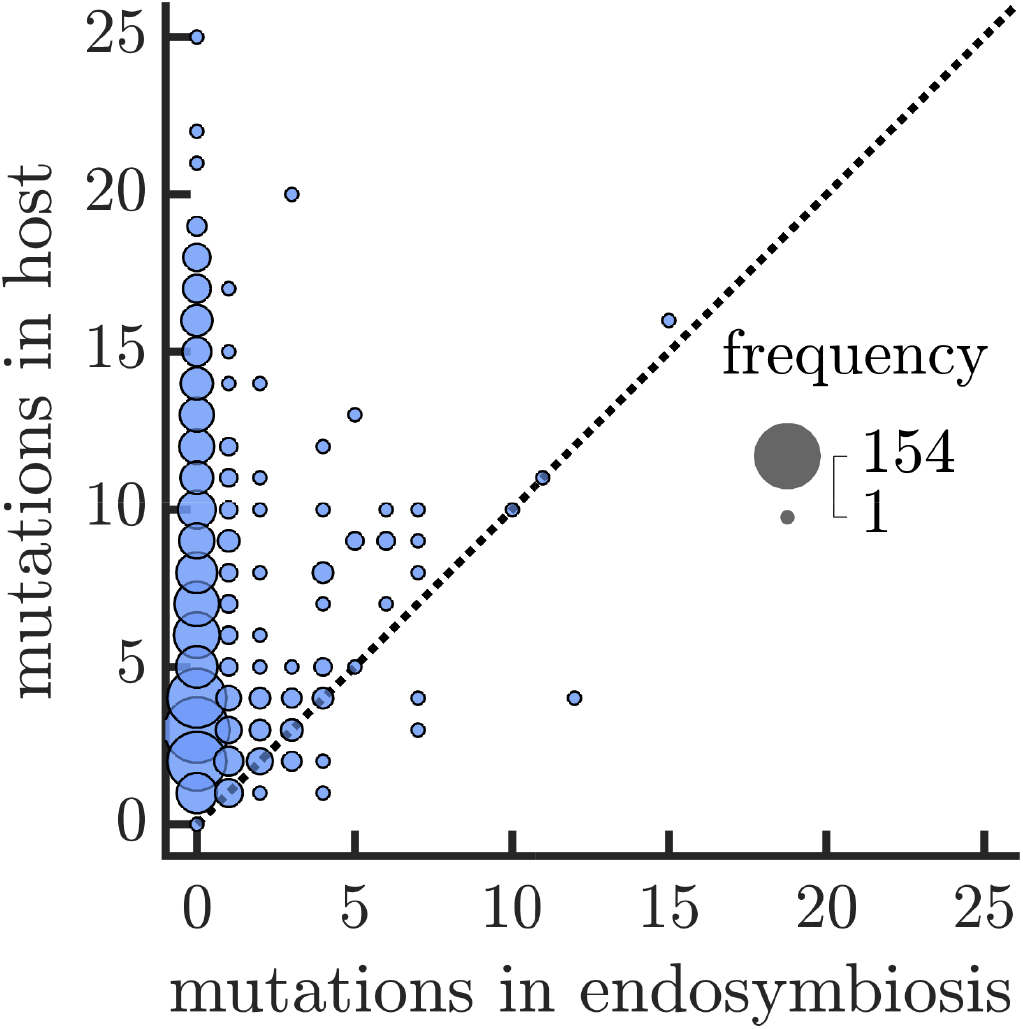
Companion to Figure 6C. The plot is the same as in Figure 6C except instead the data is from the AGORA database instead of the CarveMe database.

**Figure S12:**
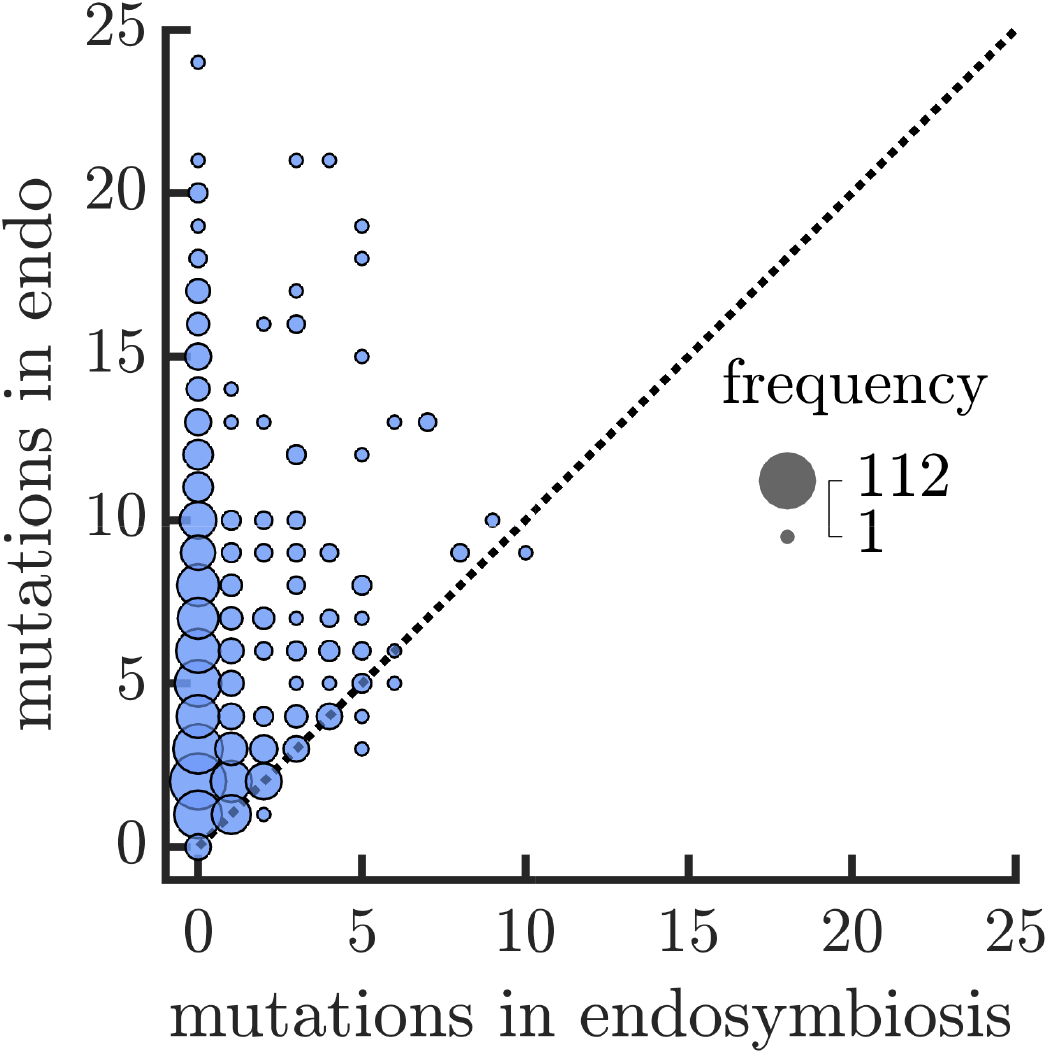
Companion to Figure 6D. The plot is the same as in Figure 6D except instead the data is from the AGORA database instead of the CarveMe database.

